# Usutu virus African 3.1 lineage, Portugal, 2021-2023

**DOI:** 10.1101/2024.12.04.626753

**Authors:** João Queirós, Tatiana Silva, Catarina Fontoura-Gonçalves, Inês Magalhães, Alberto Moraga, Marinela Contreras, Tereza Almeida, Ana M. Lopes, Joana Abrantes, Luís P. da Silva, Marisa Rodrigues, João Basso Costa, Gonçalo de Mello, David Gonçalves, Paulo Célio Alves, Ursula Höfle

## Abstract

**Background:** Usutu virus (*Orthoflavivirus usutuense*, USUV), a neurotropic arthropod-borne RNA virus of the family Flaviviridae, is a zoonotic virus that has spread throughout the European continent over the last three decades, since its emergence in Italy in 1996. However, no cases of USUV have been reported in Portugal so far.

**Material and methods:** In the scope of an active surveillance program for *Orthoflavivirus*, we collected growing feather samples from 249 red-legged partridges (*Alectoris rufa*) hunted in southern Portugal during the 2021-2023 hunting seasons. Samples positive for USUV were subjected to whole genome sequencing and strain characterization.

**Results:** Two partridges tested positive for USUV. Phylogenetic analyses of whole and partial genomes assigned the USUV strains to the African 3 lineage, specifically the African 3.1 sub-lineage.

**Conclusions:** Our study confirms, for the first time, the circulation of USUV in wild birds in Portugal. Active surveillance of hunted partridges proved to be a useful, accessible, and cost-effective method for USUV monitoring, further supporting their value as effective sentinels for *Orthoflavivirus* surveillance. Given the ongoing circulation of USUV and the increasing risk of its spillover to other domestic and wild animals, and humans, additional efforts are needed to improve virus surveillance in Portugal from a One Health perspective.

## Introduction

Emerging pathogens, particularly those transmitted by vectors, have been altering their dynamics due to global environmental changes that affect vector distributions and allow pathogens to spread to and persist in new areas (1). A notable example is the Usutu virus (*Orthoflavivirus usutuense*, USUV), a neurotropic, zoonotic virus in the genus *Orthoflavivirus* of the family Flaviviridae. Part of the Japanese encephalitis virus serocomplex, USUV was first identified in South Africa in 1959 and emerged in Europe in 1996 (2). Since then, it has rapidly spread across the European continent to at least 21 countries (Table S1, Supplementary data).

Like other orthoflaviviruses, USUV has an enzootic cycle, with birds as amplifying hosts and mosquitoes as vectors (3,4). Birds have been associated with multiple virus introductions during migrations, although the virus is endemic in some regions (4). USUV has been isolated mainly from birds (54.8% of the isolates from at least 53 species) and mosquitoes (33.1% from at least 16 species), but also from humans (4.1%) and other mammals 0.6% from at least four species), and ticks (0.4% from at least two species, Table S2, Supplementary data). Serological evidence also suggests virus circulation in other mammal species (5).

Phylogenetic studies of isolated strains have identified eight evolutionary lineages, five European and three African, often circulating simultaneously in Europe (3) (Figure 1). Notably, full genome analysis allowed the identification of three sub-lineages within the African 3 lineage, with the 3.1 sub-lineage being identified across Western Europe, from Germany to Spain (6).

**Figure 1.**
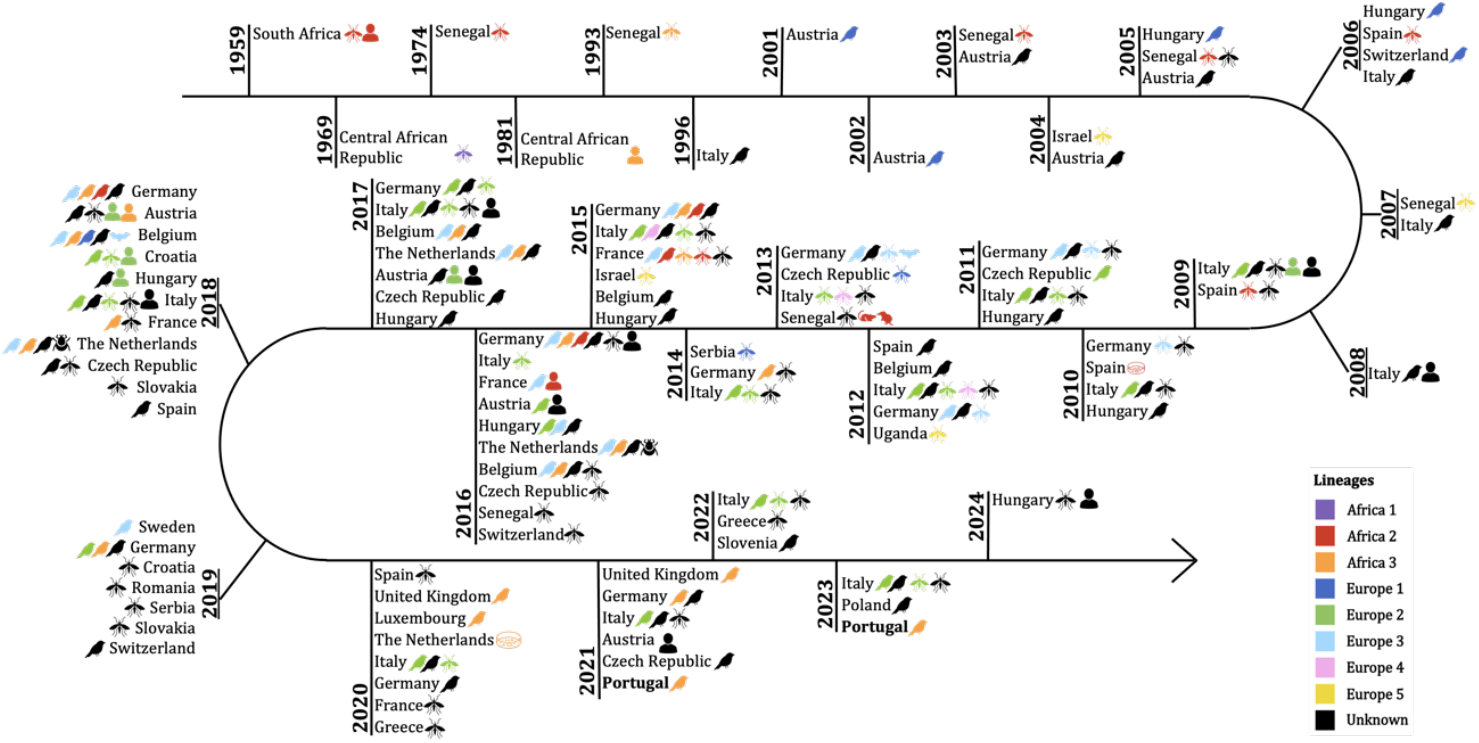
Timeline of USUV cases reported in the literature, since its emergence in 1959 to 5 November 2024. This also includes the two USUV cases described herein in Portugal, in 2021 and 2023 (bold). USUV lineages are classified based on genomes. Additional information can be consulted in Table S4, Supplementary data.

In Portugal, however, no cases of USUV have been reported so far. Following the outbreak of Bagaza virus in red-legged partridges (*Alectoris rufa*) in southern Portugal in 2021 (7), an active surveillance program for *Orthoflavivirus* in hunted partridges was implemented to identify the circulating viruses and to characterize their strains. The red-legged partridge is among the most important game birds in South-Western Europe, with millions hunted each year (8). It belongs to the family Phasianidae and has previously been suggested as a good sentinel species for *Orthoflavivirus* surveillance based on serological data (9).

## Material and methods

### Study area and sampling

The study was carried out in southern Portugal, in hunting grounds located in the municipalities of Mértola and Serpa (Figure S1, Supplementary data). This region is characterized by a Mediterranean climate and vegetation, and a sloping terrain, where active landscape management is mainly focused on improving the habitat suitability for the red-legged partridge, including cultivated parcels interspersed with natural shrubs and cork oak (*Quercus suber*) and holm oak (*Quercus ilex*) trees, which promote a high abundance of native wild populations. The hunting season for this species is between October and December and thousands of birds are shot each year. We took advantage of the regular hunting season to collect growing feather samples from 249 apparently healthy specimens between October 2021 and December 2023 (Table S3). Hunting activities were conducted independently of this study. Feathers were preserved in RNAlater at -80º C until lab processing.

### RNA extraction and RT-qPCR assays

Total RNA was extracted from the pulp of growing feathers using a MagMAX CORE Nucleic Acid Purification kit™ in the KingFisher Apex instrument™. Samples were screened for avian orthoflaviviruses using a duplex quantitative reverse transcription PCR (RT-qPCR) for the simultaneous and differential detection of Japanese encephalitis (JE) and Ntaya *Orthoflavivirus* serocomplexes (10). Positive samples to the JE serocomplex were further screened for detection of USUV using an RT-qPCR for amplification of part of the nonstructural protein 5 (NS5) gene region of USUV (2).

### RNA virus library preparation and sequencing

The positive samples to USUV were used for RNA library preparation. RNA concentration and quality were assessed with a Qubit 4™ Fluorometer (RNA HS Assay Kit™) for precise quantification, and purity was confirmed via NanoDrop™ Lite Spectrophotometer (Thermo Fisher Scientific, MA, USA), examining A260/A280 and A260/A230 ratios. Residual genomic DNA was removed by DNase I treatment (NZYtech, Portugal), and RNA was further purified using the GeneJET RNA Purification Kit (Thermo Scientific, MA, USA). RNA integrity and concentration were re-evaluated on an Agilent 4200 TapeStation, with RNA values measured per the manufacturer’s guidelines. Ribosomal RNA was removed using the NEBNext rRNA Depletion Kit v2 (Human/Mouse/Rat; New England Biolabs), and the rRNA-depleted RNA was used to prepare libraries with the NEBNext Ultra II Directional RNA Library Prep Kit for Illumina (New England Biolabs) following the manufacturer’s protocol. Sequencing was conducted on a NovaSeq 6000 platform (150 bp paired-end) at Macrogen (Seoul, Korea).

### Phylogenetic analysis

The genome consensus sequences of the two positive partridges were constructed using Geneious Prime® 2024.0.3 (https://www.geneious.com/biopharma). Paired-end data were combined for each sample and then cleaned using Bbulk plug-in and merged using BBmerge. Sequences were then mapped against a USUV reference genome (NCBI reference sequence: NC_006551.1) using default features with medium/low sensitivity 5x and no trim function.

All publicly available sequences under the terms “*Orthoflavivirus usutuense*” and “Usutu virus” were retrieved from the online database NCBI Virus (https://www.ncbi.nlm.nih.gov/labs/virus/) on 5 November 2024 (n = 1392, Table S4) and downloaded into Geneious 9.2.4 (https://www.geneious.com). Unique genomes (n = 675) were aligned in MAFFT and cropped to include only the known coding region (10 305 bp), as in the reference sequence genome NC_006551.1. Redundant sequences and sequences containing ambiguities (Ns) were removed from the alignment, resulting in a total of 537 single coding region genomes, which were used for further analyses. ModelFinder from IQTree was used to select the best-fitting model according to BIC (11). Maximum likelihood phylogenetic tree was constructed with IQ-TREE web server (12) using GTR+F+I+G4 and 1 000 replications of ultrafast bootstrap resampling (13) and SH-aLRT test (14). The consensus tree was visualized in iTOL (15) and further edited in Inkscape v1.3.2 (https://inkscape.org/release/inkscape-1.3.2/).

## Results and Discussion

Two out of the 249 partridges tested positive for USUV (0.8%), *Alectoris rufa*-1 (a juvenile female hunted in Serpa in November 2021; cycle threshold 32.16) and *Alectoris rufa*-2 (an adult female hunted in Mértola in November 2023; cycle threshold 21.26). This confirms previous serological evidence of USUV circulation in southern Portugal (8).

In the high-throughput sequencing analysis, we obtained 558,636 total reads for *Alectoris rufa*-1 (Serpa, 2021) and 29,220,938 total reads for *Alectoris rufa*-2 (Mértola, 2023). For *Alectoris rufa*-1, 113 paired reads were mapped to the reference genome, forming 10 contig fragments of a total of 1,401 base pairs with a minimum coverage of 5 bp. These included positions 298-510 (213bp), 964-1,065 (102bp), 1,474-1,572 (99bp), 2,308-2,436 (129bp), 3,292-3,477 (186bp), 4,864-4,974 (111bp), 5,626-5,742 (117bp), 6,661-6,750 (90bp), 7,336-7,512 (177bp) and 7,801-7,977 (177bp) of reference sequence genome (NC_06551.1) (Accession numbers: PQ672170-79). For *Alectoris rufa*-2, 7,550 paired reads were mapped to the USUV reference genome, forming a contig of a total of 11,108 base pairs with an average coverage of 105x (minimum coverage of 14x and maximum of 307x) (Accession number: PQ677887). Pairwise distance analysis revealed a high degree of similarity (pairwise identity of 99.4%) between the protein coding region of *Alectoris rufa*-1 and the corresponding region in the complete genome of *Alectoris rufa*-2.

Phylogenetic analyses of partial (*Alectoris rufa*-1) and complete (*Alectoris rufa-*2) genomes assigned the USUV strains to the African 3 lineage (Figure 2). First described in the Central African Republic in 1981 (human case), and then in Senegal in 1993 (*Culex univittatus*) and 2007 (*Culex neavei*), this lineage emerged in Europe in 2014, in a *Turdus merula* found dead in Germany (Figure 1). Since then, it has spread throughout Western Europe, and three sub-lineages now co-circulate in some countries (6). We detected the African 3.1 sub-lineage (Figure 2a), initially found in Germany in 2016, in Belgium, France, and the Netherlands in 2018, and in Luxembourg and Spain in 2020 (Figure 2b). The widespread distribution of the African 3.1 sub-lineage and other USUV (sub)lineages in wild birds and mosquitoes in Western European countries suggests the temporal persistence and evolution of the virus in the continent.

**Figure 2.**
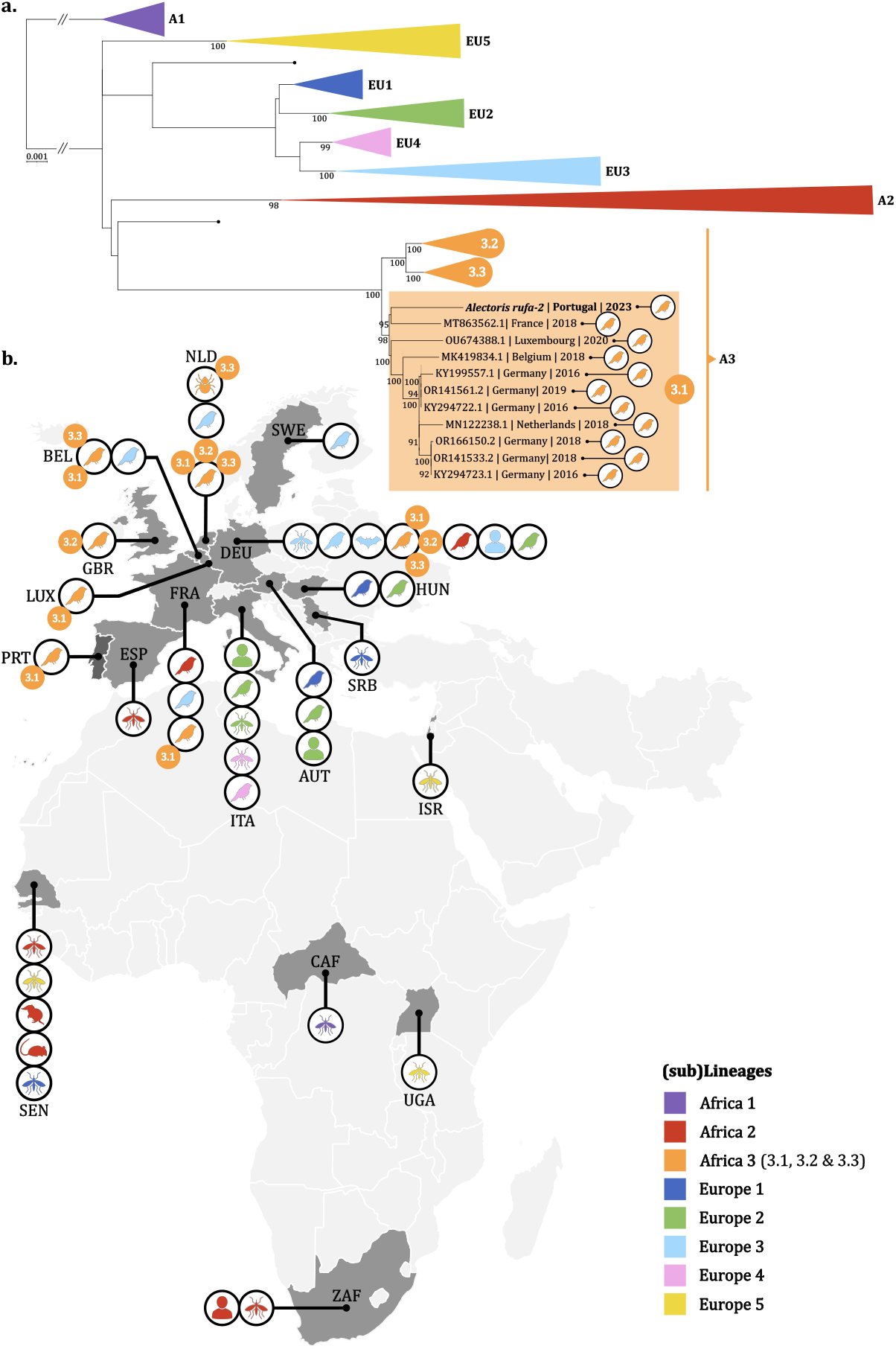
Whole genome USUV data available in the NCBI Virus Database on 5 November 2024. **a.** Maximum likelihood phylogenetic tree constructed using the virus isolated from the *Alectoris-rufa*-2 and 537 unique complete coding region sequences available. USUV lineages are represented by color: purple, Africa 1 (A1); red, Africa 2 (A2); orange, Africa 3 (A3); dark blue, Europe 1 (EU1); green, Europe 2 (EU2); light blue, Europe 3 (EU3); pink, Europe 4 (EU4); and yellow, Europe 5 (EU5). Sub-lineages are shown for the African 3 lineage, and accession numbers and hosts are depicted only for the African 3.1 sub-lineage, defined according to (6) (see details of the list of genomes included in the analyses in Table S4, Supplementary data); **b**. Geographic distribution of USUV genomes without ambiguities (n = 646) by lineage and host in which they were isolated. Countries are identified by 3-letter International Organization for Standardization codes (https://www.iso.org). USUV sub-lineages are shown for the African 3 lineage only. Map created using qGIS Desktop 3.34.3 (http://www.qgis.org) and open data from Free Vector Maps (https://freevectormaps.com).

This might represent an increasing risk of USUV spillover to other domestic and wild animals, as well as humans (16). Additional efforts to improve disease surveillance in wildlife, mosquitoes, domestic animals, and humans are required, especially in Portugal, where no USUV-related outbreaks or deaths have been reported, despite evidence of its circulation since 2018 (9).

Our study shows that vascular growing feathers from apparently healthy wild partridges hunted during the regular hunting season is a useful, accessible, and cost-effective sampling approach for active surveillance of USUV and its genomic characterization. Despite the temporal limitation to the hunting season, this approach has a wide potential application in Portugal and abroad. Red-legged partridges are widely distributed and hunted across Western Europe, from introduced populations in the United Kingdom to native populations in France, Spain, Italy and Portugal (8). This is the second USUV case detected in red-legged partridges (Table S2, Appendix) and our results provide additional support for considering this species as a good sentinel for *Orthoflavivirus* surveillance (9).

## Conclusion

Our study detects USUV African 3.1. sub-lineage in red-legged partridges hunted in southern Portugal in November 2021 and 2023, confirming previous serological evidence of USUV circulation in wild birds in the country. Active surveillance through the collection of vascular growing feathers from apparently healthy wild partridges hunted during the regular hunting season proved to be a useful, accessible, and cost-effective method for USUV monitoring. This reinforces previous evidence supporting red-legged partridges as a good sentinel for *Orthoflavivirus* surveillance. Given the ongoing circulation of USUV and the increasing risk of its spillover to other domestic and wild animals, and humans, additional efforts are needed to improve virus surveillance in Portugal from a One Health perspective.

## Acknowledgements

We are grateful for the support of Herdade de Vale de Perditos and Herdade das Romeiras, and their staff for their help collecting the data.

