## Supplementary data for "Usutu virus African 3.1 lineage, Portugal, 2021-2023"

### Supplementary Tables

Table S1……………………………………………………………………………….2

Table S2……………………………………………………………………………….3

Table S3……………………………………………………………………………….4

Table S4……………………………………………………………………………….4

### Supplementary Figures

Figure S1…………………………………………………………………………...….4

**References**………………………………………………………………………….7

### Supplementary Tables

**Table S1**. Number of Usutu virus sequences reported in the National Center for Biotechnology Information (NCBI) database on 5th November 2024 by country.

| Continent | Country | NCBI sequences |
|---|---|---|
| **Europe** | Austria | 83 |
| | Belgium | 46 |
| | Croatia | 4 |
| | Czech Republic | 40 |
| | France | 14 |
| | Germany | 410 |
| | Greece | 3 |
| | Hungary | 22 |
| | Israel | 9 |
| | Italy | 468 |
| | Luxembourg | 1 |
| | Netherlands | 144 |
| | Poland | 34 |
| | Romania | 1 |
| | Serbia | 2 |
| | Slovakia | 16 |
| | Slovenia | 1 |
| | Spain | 10 |
| | Sweden | 1 |
| | Switzerland | 49 |
| | United Kingdom | 2 |
| | Total | 1282 |
| **Africa** | Central African Republic | 2 |
| | Senegal | 19 |
| | Uganda | 2 |
| | South Africa | 4 |
| | Total | 27 |
| **Unknown** | | 5 |
| **Total** | | 1392 |

**Table S2.** Number of Usutu virus sequences reported in the National Center for Biotechnology Information (NCBI) database on 5th November 2024 by taxonomic group.

| Taxonomic group | Species/genus/family | NCBI sequences |
|---|---|---|
| Bird | *Acanthis flammea* | 2 |
| | *Actitis hypoleucos* | 1 |
| | *Alauda arvensis* | 3 |
| | *Alectoris rufa* | 1 |
| | *Alopochen aegyptiaca* | 1 |
| | *Anas platyrhynchos* | 5 |
| | *Anatidae* | 1 |
| | *Anser indicus* | 2 |
| | *Asio otus* | 4 |
| | *Aves* | 1 |
| | *Branta ruficollis* | 3 |
| | *Bubo scandiacus* | 2 |
| | *Caprimulgus europaeus* | 2 |
| | *Ciconia ciconia* | 1 |
| | *Columba livia* | 2 |
| | *Columba palumbus* | 2 |
| | *Columba sp.* | 1 |
| | *Columbidae* | 1 |
| | *Corvus cornix* | 1 |
| | *Cyanistes caeruleus* | 4 |
| | *Cygnus olor* | 1 |
| | *Erithacus rubecula* | 1 |
| | *Erythrura prasina* | 1 |
| | *Falco subbuteo* | 1 |
| | *Fringilla coelebs* | 4 |
| | *Garrulus glandarius* | 4 |
| | *Gracula religiosa* | 1 |
| | *Laridae sp.* | 1 |
| | *Larus crassirostris* | 2 |
| | *Larus michahellis* | 2 |
| | *Leucopsar rothschildi* | 1 |
| | *Melanitta nigra* | 1 |
| | *Mergus squamatus* | 1 |
| | *Motacilla alba* | 2 |
| | *Muscicapa striata* | 1 |
| | *Oeciacus hirundinis* | 1 |
| | *Panurus biarmicus* | 1 |
| | *Parus major* | 3 |
| | *Passer domesticus* | 5 |
| | *Passer montanus* | 1 |
| | *Phasianus colchicus* | 1 |
| | *Phoenicurus phoenicurus* | 1 |
| | *Pica pica* | 2 |
| | *Pyrrhula erythaca* | 1 |
| | *Pyrrhula pyrrhula* | 2 |
| | *Regulus regulus* | 10 |
| | *Serinus canaria* | 3 |
| | *Sitta europaea* | 2 |
| | *Streptopelia decaocto* | 1 |
| | *Strix aluco* | 1 |
| | *Strix nebulosa* | 53 |
| | *Sturnus vulgaris* | 7 |
| | *Surnia ulula* | 3 |
| | *Sylvia atricapilla* | 3 |
| | *Tachyeres sp.* | 2 |
| | *Troglodytes troglodytes* | 1 |
| | *Turdidae* | 2 |
| | *Turdus iliacus* | 3 |
| | *Turdus merula* | 525 |
| | *Turdus philomelos* | 12 |
| | *Turdus pilaris* | 3 |
| | Unknown | 44 |
| | Total | 757 |
| Human | Homo sapiens | 57 |
| Mammal | *Crocidura sp.* | 1 |
| | *Mastomys natalensis* | 3 |
| | *Pipistrellus pipistrellus* | 3 |
| | *Rattus rattus* | 1 |
| | Total | 8 |
| Mosquito | *Aedes albopictus* | 2 |
| | *Aedes japonicus* | 1 |
| | *Aedes vexans* | 1 |
| | *Anopheles claviger* | 1 |
| | *Anopheles maculipennis* | 1 |
| | *Culex sp.* | 24 |
| | *Culex modestus* | 6 |
| | *Culex neavei* | 12 |
| | *Culex perexiguus* | 7 |
| | *Culex perfuscus* | 2 |
| | *Culex pipiens* | 377 |
| | *Culex pipiens complex* | 8 |
| | *Culex pipiens pipiens* | 2 |
| | *Culex pipiens sensu lato* | 11 |
| | *Culex poicilipes* | 1 |
| | *Culex torrentium* | 1 |
| | *Culex univittatus* | 3 |
| | *Culicidae* | 1 |
| | Total | 461 |
| Ticks | *Ixodes frontalis* | 1 |
| | *Ixodes ricinus* | 3 |
| | *Ixodida* | 1 |
| | Total | 5 |
| Vero cell | | 2 |
| Unkown | Total | 102 |
| **Total** | | 1392 |

**Table S3**. Number of hunted red-legged partridges analyzed in the municipalities of Mértola and Serpa between October 2021 and December 2023.

| Municipalities | 2021 | 2022 | 2023 | | | Total |
|---|---|---|---|---|---|---|
| | November | November | October | November | December | |
| Mértola | 0 | 0 | 40 | 30 | 0 | 70 |
| Serpa | 46 | 80 | 0 | 29 | 24 | 179 |
| Total | 46 | 80 | 40 | 59 | 24 | 249 |

**Table S4**. Excel file with Usutu virus sequences available in the National Center for Biotechnology Information (NCBI) database on 5 November 2024 (see Excel File attached).

**Supplementary Figures**

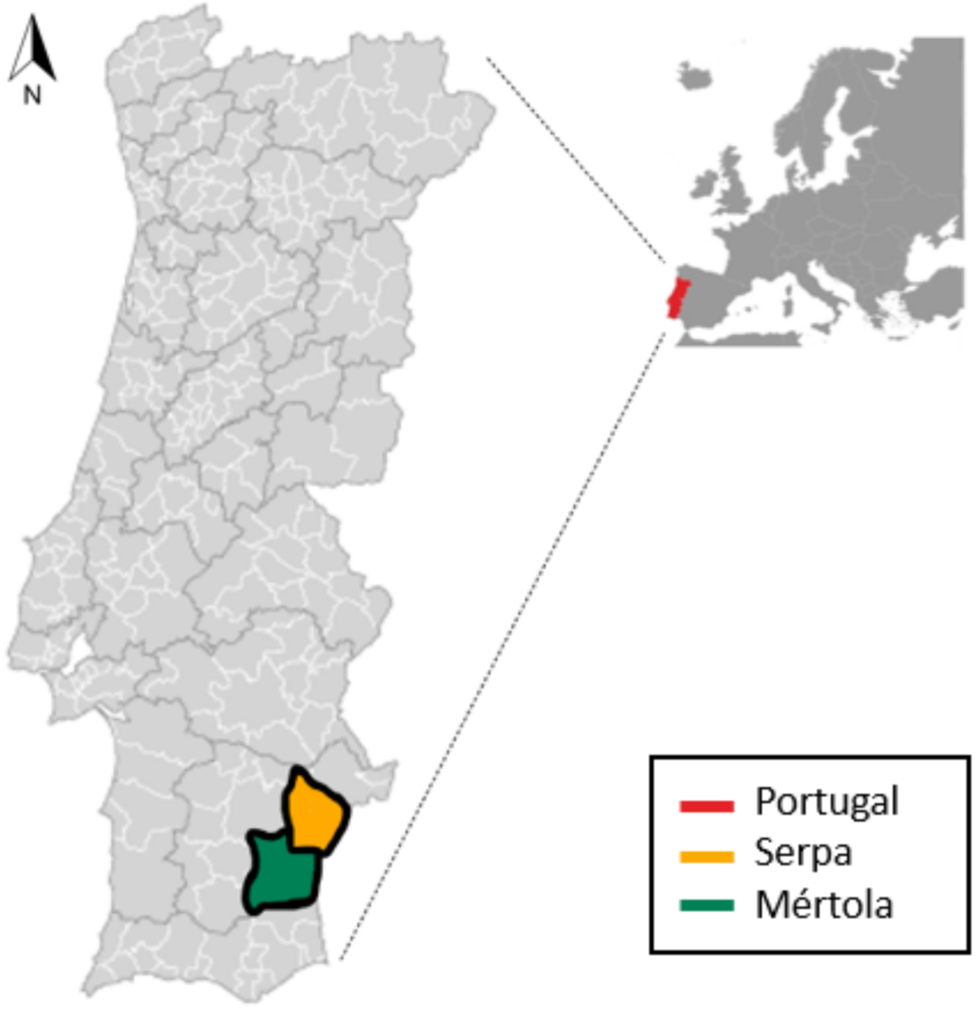


**Figure S1**. Study area and sampled municipalities of Mértola and Serpa in Southern Portugal.
